# High population density limits predator access in Antarctic fur seal breeding colonies

**DOI:** 10.64898/2026.04.07.716769

**Authors:** Johannes Bartl, Ane Liv Berthelsen, Alexander Winterl, Cameron Fox-Clarke, Jaume Forcada, Rebecca Nagel, Joseph I. Hoffman, Ben Fabry

## Abstract

Population density can influence individual predation risk in colonial breeders through shared vigilance and predator deterrence. We investigated how predator-prey interactions are shaped by population density at two Antarctic fur seal (*Arctocephalus gazella*) breeding colonies at Bird Island, South Georgia, which differ four-fold in seal density. By deploying autonomous time-lapse cameras, we captured high-resolution images at one-minute intervals throughout the breeding season. Using a YOLOv8 neural network, we identified fur seal adult males, females and pups, as well as three predator-scavenger bird species: giant petrels (*Macronectes spp*.), brown skuas (*Stercorarius antarcticus*) and snowy sheathbills (*Chionis alba*). Abundance patterns corresponded to the known foraging and breeding behaviours of these species. Differences in seal density between the colonies were mainly driven by adult females and their pups, but not adult males. The ratios of predatory birds to pups were markedly lower at the high-density colony, while scavenger to pup ratios remained similar. Spatial analyses revealed that predators were largely excluded from areas of high seal density, whereas scavengers overlapped extensively with pups in both colonies. This study demonstrates the value of remote observation in resolving predator-prey interactions and illustrates how density can shape predation risk in a colonial breeder.

## Background

Population density can strongly influence predation risk, particularly in colonial breeders (Hamilton 1971; Kramer *et al*. 2009; Morrell & James 2008). In breeding colonies, predation risk may be reduced through shared vigilance (Beauchamp 2008; Hamilton 1971; Pulliam 1973), which increases the likelihood of detecting predators. Predation risk may also be reduced as a consequence of enhanced predator deterrence or even exclusion through collective anti-predator behaviours (Guidos *et al*. 2023; Kazama & Watanuki 2010), and by the dilution effect (Bednekoff & Lima 1998), whereby individual risk decreases as group size increases. These processes are often framed within the context of an Allee effect (Allee 1931), where individual fitness increases with population density.

Allee effects are generally difficult to demonstrate in wild populations (Courchamp *et al*. 1999), but colonial breeding systems provide a setting in which fitness variation can be studied in relation to natural variation in density over space and time (Kramer *et al*. 2009). One such ‘natural experiment’ is provided by Antarctic fur seals (*Arctocephalus gazella*) breeding at Bird Island, South Georgia. Here, empirical evidence for a component Allee effect on pup survival was obtained by comparing past and present drivers of pup mortality (Nagel *et al*. 2021) across two neighbouring breeding colonies – the special study beach (SSB) and freshwater beach (FWB) – which differ four-fold in seal density (Meise *et al*. 2016). Historically, pup mortality was highest at SSB (the high-density colony) due to injuries from territorial males and starvation caused by mother-pup separation (Doidge *et al*. 1984; Reid & Forcada 2005). However, following a recent decline in seal abundance linked to changes in food availability due to climate change (Forcada *et al*. 2023; Forcada & Hoffman 2014), pup mortality is now higher at FWB (the low-density colony), largely due to predation (Nagel *et al*. 2021).

The main terrestrial predators of Antarctic fur seal pups are the northern and southern giant petrels (*Macronectes halli* and *M. giganteus*, Nagel et al. 2022). These large seabirds feed mainly on fur seal placentae and carcasses as well as penguin carcasses (Hunter 1983) during the austral summer (November-February). Giant petrels attack pups by pecking and drowning (Nagel *et al*. 2022), targeting small, unattended pups in areas with a low density of adult seals (Nagel *et al*. 2022). The brown skua (*Stercorarius antarcticus*) feeds almost exclusively on fur seal placentae and carrion during their incubation period, which overlaps with the fur seal pupping season (Phillips *et al*. 2004) and has also been anecdotally observed opportunistically preying on pups (Nagel *et al*. 2022). Snowy sheathbills (*Chionis alba*) are smaller opportunistic predator-scavengers with a flexible diet including sea algae, crustaceans and marine mammal placentae and carcasses (Favero 1996). These bird species are all closely associated with fur seal breeding colonies and together form a predator to scavenger gradient.

The breeding season of Antarctic fur seals is concentrated within the austral summer. Adult males arrive at breeding colonies in early November, where they establish territories (McCann 1980). About a month later, the adult females come ashore to give birth to pups conceived during the previous breeding season (Duck 1990). Parturition generally takes place within one to two days, and females enter oestrus around seven days afterwards (Duck 1990; Lunn *et al*. 1994). After mating, the females alternate nursing their pups ashore with foraging trips at sea (Forcada & Staniland 2009; Lunn *et al*. 1994). During this period, the breeding colonies become important foraging sites for predatory and scavenging birds.

During early development, Antarctic fur seal pups are largely confined to their natal breeding colony. Although their mothers protect them from predators (Nagel *et al*. 2021) the pups are left unattended while their mothers forage at sea (Lunn *et al*. 1994), increasing their risk of predation (Nagel *et al*. 2021). Consequently, predation risk is expected to be determined by the spatio-temporal overlap between pups and predatory birds in the absence of adult females. However, investigating fine-scale predator-prey interactions remains challenging, as traditional observation methods often capture only a small fraction of predation events. Nevertheless, recent advances in automated monitoring using fixed cameras combined with automatic detection algorithms now allow for a continuous observation of breeding colonies, providing a non-invasive and labour-efficient approach for investigating spatiotemporal abundance patterns and predator-prey interactions (Berthelsen & Bartl 2026).

To investigate these dynamics, we deployed time-lapse cameras at FWB and SSB, which recorded images at 1-minute intervals over two months, producing 66,645 and 61,569 images respectively. Following the approach introduced in (Berthelsen & Bartl. 2026) a custom-trained neural network was used to identify and locate animals within the images from FWB, and the resulting dataset is reused here. In the present study, we applied a similar approach at SSB. This allowed us to directly compare predator-prey patterns between the two colonies and to evaluate how well the methodology transfers between colonies differing in size, density and visibility. By comparing model performance and temporal abundance patterns of Antarctic fur seals and their associated bird species, we evaluated the robustness and generalisability of this approach for seal colony monitoring.

Using the neural network annotated datasets, we investigated the temporal and spatial abundance of Antarctic fur seals and three associated bird species. Given the lower predation risk previously recorded at SSB (Nagel *et al*. 2021), we hypothesised that (i) predatory bird abundance would be lower at the high-density colony, and (ii) that this difference would be reflected in bird-to-pup abundance ratios, as fewer birds per pup should reduce the likelihood of predator encounters, and thus predation risk. The contrast in seal densities between the two colonies was described a decade ago (Meise *et al*. 2016), but whether these differences reflect variation between sexes or life history stages remain unclear. We therefore quantified colony-specific densities of adult males, adult females and pups, hypothesising that (iii) the overall differences in seal density reflected differences between these groups. Finally, we analysed the extent of spatial overlap between birds and pups, assuming that spatial overlap serves as a proxy for predation risk, and that adult females actively protect their pups against predators (Nagel *et al*. 2021). We hypothesised that (iv) the higher density of SSB would result in the spatial exclusion of predatory but not scavenging bird species.

## Methods

### (a) Study sites

This study was conducted at Bird Island, South Georgia (54°00’24.8” S, 38°03’04.1” W) during the austral summer of 2020–2021 at two Antarctic fur seal breeding colonies (Figure 1). Freshwater Beach (FWB) lies directly in front of the British Antarctic Survey (BAS) field base. This site is characterized by an open beach intersected by a shallow stream that forms natural corridors and influences the spatial distribution of seals within the colony (Berthelsen & Bartl 2026). Approximately 200 metres south of FWB lies the Special Study Beach (SSB), a small cobblestone breeding beach separated from adjacent breeding sites by a cliff on the east side, open sea on the west and rocky ridges to the north and south. Here, a scaffold walkway (Doidge *et al*. 1984) provides safe access to the animals while minimising disturbance. During the breeding season, a daily census quantified the abundance of adult males, adult females and newborn pups (further details in supplementary).

**Figure 1:**
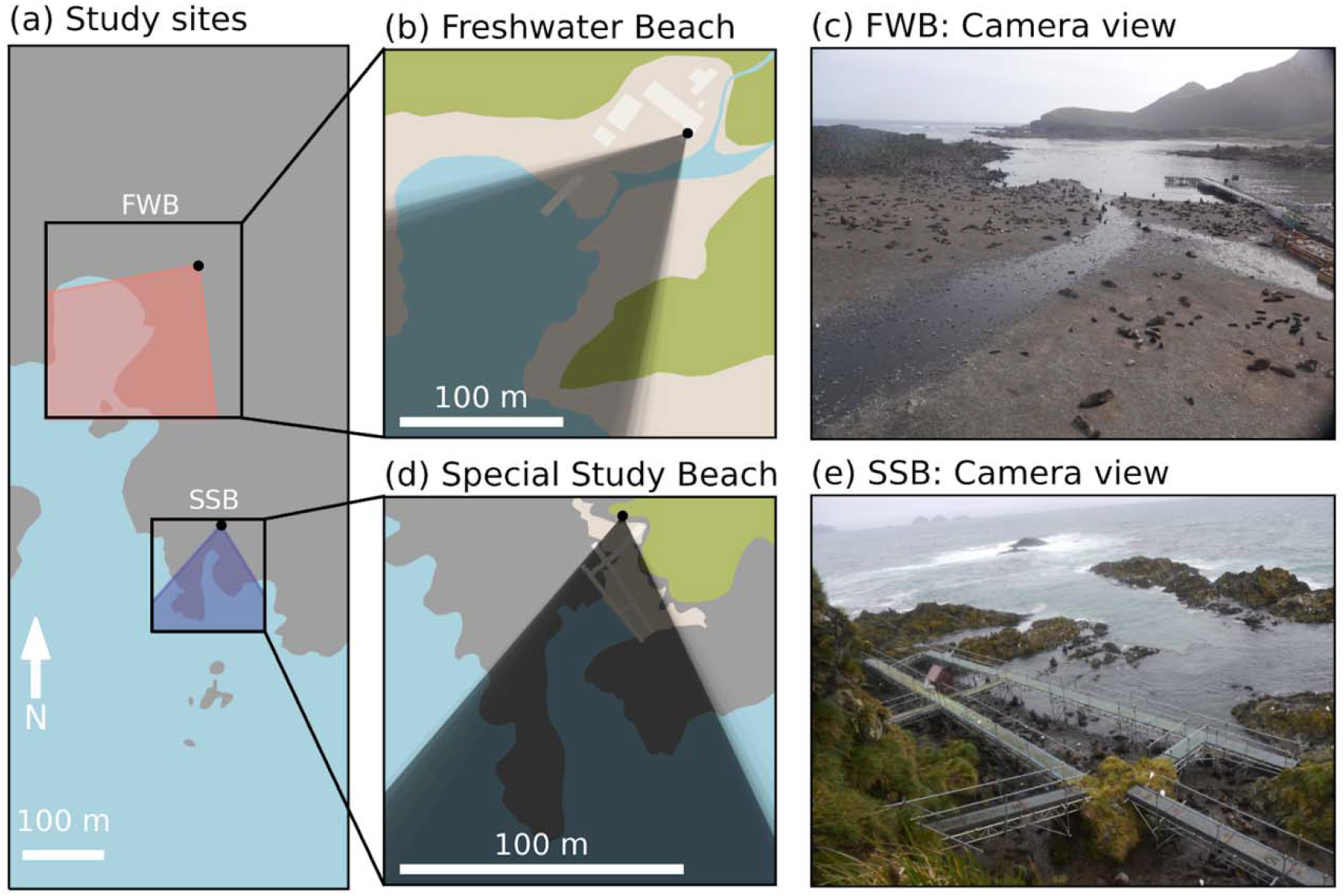
Study location and camera view. (a) Locations of the two Antarctic fur seal breeding colonies: Freshwater Beach (FWB) and Special Study Beach (SSB) at Bird Island, South Georgia. The black boxes outline the study colonies, with the total camera field of view (FOV) across the season indicated in salmon (FWB) and indigo (SSB). (b, d) Schematic illustration of each site showing the FOV used during the study; darker shading indicates areas monitored more frequently. (c, e) Example images captured by the fixed position cameras at FWB and SSB, respectively. Black points indicate the camera positions.

### (b) Remote observatories

The spatial distribution of individuals was recorded at a rate of one image per minute using an automated camera system (Winterl *et al*. 2020). Each system consisted of a 16-megapixel digital mirrorless camera (Panasonic Lumix DMC-G5) with a 14 mm lens (horizontal field of view = 23°). The cameras’ controller unit was based on an Arduino microcontroller. The camera and controller were protected in a water-resistant housing and mounted on a tripod. A 12 V car battery served as a power supply. At FWB, the camera was mounted on the field station’s tower at a height of eight meters. It recorded continuously for 59 days between 28 October and 25 December 2020, producing a total of 66,645 images. An identical camera system was installed at SSB on a tripod positioned on an elevated cliff overlooking the colony. It recorded continuously for 55 days between November 8th, 2020, and January 1st, 2021, producing 61,569 images.

### (c) Training dataset and data filtering

We randomly selected two images per day for manual annotation, resulting in 110 full-sized images. This sampling strategy ensured that the training set captured natural variation in the recording conditions, spanning diverse ambient light levels, visibility, shadows and seal densities. We annotated all animals using the image annotation tool Clickpoints (Gerum *et al*. 2017). For each observed animal, we assigned a class – taxa, species or further differentiating category – and drew a bounding box enclosing the entire animal. If an individual was partially occluded, we annotated its full extrapolated (expected) outline. We defined nine distinct classes to annotate all relevant animals on the beach. These included: elephant seal (*Mirounga leonina*), leopard seal (*Hydrurga leptonyx*), and three Antarctic fur seal categories: adult males, adult females and pups, king penguin (*Aptenodytes patagonicus*), giant petrel, brown skua and snowy sheathbill. Of these, we focused our study on the three Antarctic fur seal categories and their associated predator-scavenger bird species: giant petrel, brown skua and snowy sheathbill.

To ensure sufficient daylight and optimal visibility, we restricted the dataset to images captured between 09:00 and 17:00 local time. To systematically distinguish clear images from those obscured by environmental factors such as fog or water droplets on the lens, we established a quality baseline. To this end, we randomly selected 200 images from SSB and manually categorized these as being either suitable or unsuitable for analysis. For this subset of images, we computed the Laplacian variance (Pech-Pacheco *et al*. 2000) and determined the threshold that minimized the number of incorrectly classified images. A variance threshold of 25 proved optimal for separating the categories and was subsequently applied to filter the full SSB image set, yielding a total of 9,940 high quality images (16%) captured across 45 days.

### (d) Automated detection

For the automated analysis of the filtered dataset, we trained a neural network using the YOLOv8 architecture (Jocher *et al*. 2023). This is a state-of-the-art object detection model designed for high efficiency and accuracy. Its architecture is composed of three main components: a specialized backbone for feature extraction; a path aggregation network (PANet) neck that merges feature maps from different scales to detect objects of varying sizes; and a decoupled head that processes object classification and bounding box regression independently. We selected the YOLOv8l (Large) variant (43.7 million parameters) for its increased depth and channel width to ensure superior feature retention and detection accuracy compared to smaller, speed-optimized versions.

To prepare the dataset of 110 full-sized annotated images for training, we applied a sliding window approach of generating 512×512 pixel crops with a 50% overlap. This resulted in 5,291 crops that were randomly split into a training set (75%) and a validation set (25%). To improve model robustness and generalization, we applied data augmentation techniques during training. These included random translations, scaling and colour adjustments. Additionally, we utilized Mosaic augmentation, which combines four training images into a single input, which enhances the model’s ability to detect objects within complex contexts. We trained the model for 150 epochs and achieved a mean Average Precision (mAP_50_) of 0.8. We optimized the confidence threshold (0.65) to maximize the F1 score. The resulting F1 score averaged over all classes was 0.77; the per class-F1 score can be seen in supplementary figure 1.

To apply the network, which was only trained on cropped images, to full sized images, we utilized the Slicing Aided Hyper Inference (SAHI) Python package (Akyon *et al*. 2022). This uses a sliding window approach with a slice size of 512×512 pixels and height and width overlap ratios of 0.2. To consolidate predictions from overlapping crops, we merged bounding boxes using Non-Maximum Suppression (NMS). This procedure yielded a total of 0.69 million detections on SSB.

### (e) Data analyses

The performance of the neural network applied to images from SSB was assessed by comparing automated counts with manually generated counts, following the same validation procedure used for the neural network developed for FWB (Berthelsen & Bartl 2026). This confirmed close agreement between the automated and manual approaches (Supplementary Figure 2 and 3). In addition, automated seal counts were compared to census counts, which demonstrated near-perfect agreement for adult males and good agreement for adult females, but lower agreement for pups mostly due to missed detections (false-negative, Supplementary Figure 4). Daily patterns of Antarctic fur seal categories and the three focal bird species were quantified throughout the season on SSB.

To characterise colony-specific patterns in seal densities, we quantified the local density (number per area) of each Antarctic fur seal category. We specified the main breeding areas at FWB and SSB by identifying the 99% occupancy area for adult females and pups using kernel density estimation. In each image, we implemented Voronoi tessellation for each seal category and all seals combined, resulting in a designated area for each individual occupying the main breeding area. We then calculated densities as the inverse of the assigned area (animals / m^2^) and visualised the resulting distributions using histograms separately for each seal category at FWB and SSB. Because adult females and pups are particularly at risk of being occluded by the scaffolding at SSB, Voronoi cells intersecting the scaffolding structure were excluded for these categories, yielding unobstructed density estimates used throughout the main analysis. The uncorrected distributions are provided for comparison in the supplementary material (Supplementary Figure 5).

To investigate the relationship between the abundance of predator-scavenger birds and Antarctic fur seal pups, we calculated bird-to-pup abundance ratios for each day following the date of the first-born pup at each colony (FWB = November 18th, SSB = November 22nd). We computed a 7-day rolling bird-to-pup ratio as a proxy for temporal variation in predation risk, under the assumption that a lower number of birds per pup reduces the likelihood of a pup encountering an avian predator. A season-integrated ratio was also calculated by dividing total bird counts by total pup counts across all days following the start of the pupping season at each colony. Differences in rolling bird-to-pup ratios between FWB and SSB were statistically evaluated for each species using two-sided Mann-Whitney U tests.

To evaluate spatial associations between Antarctic fur seal pups and the three focal bird species, and to test whether the higher seal densities at SSB lead to the exclusion of avian predators from the main breeding area, we compared their colony-level spatial distributions. For pups and the three focal bird species, we generated 99% occupancy areas using kernel density estimation separately for each colony. To quantify differences in spatial associations with pups among the bird species across colonies, we calculated the proportion of the pup occupancy area that overlapped with the respective bird occupancy area.

## Results

### (a) Temporal trends in abundance at SSB

To characterise temporal trends in abundance at SSB throughout the breeding season, we plotted the daily maximum counts of each focal class from November 9th until January 1st (*n* = 54 days, Figure 2a, b). During the first two weeks of the Antarctic fur seal breeding season, adult males predominated in the colony and gradually increased in abundance until the birth of the first pup (November 22nd). Thereafter, adult male abundance remained stable for a period of approximately three weeks, before gradually declining during the main mating period. Adult females were scarce during the first two weeks, but then increased rapidly, reaching peak abundance on December 10th, after which numbers declined. Pup abundance mostly mirrored female abundance. These temporal trends are similar to those observed at FWB (Berthelsen & Bartl 2026) and align with the known breeding phenology of Antarctic fur seals (Duck 1990). The numbers of both giant petrels and brown skuas remained low throughout the season, while snowy sheathbills were present at higher numbers, peaking between December 9th and December 25th.

**Figure 2:**
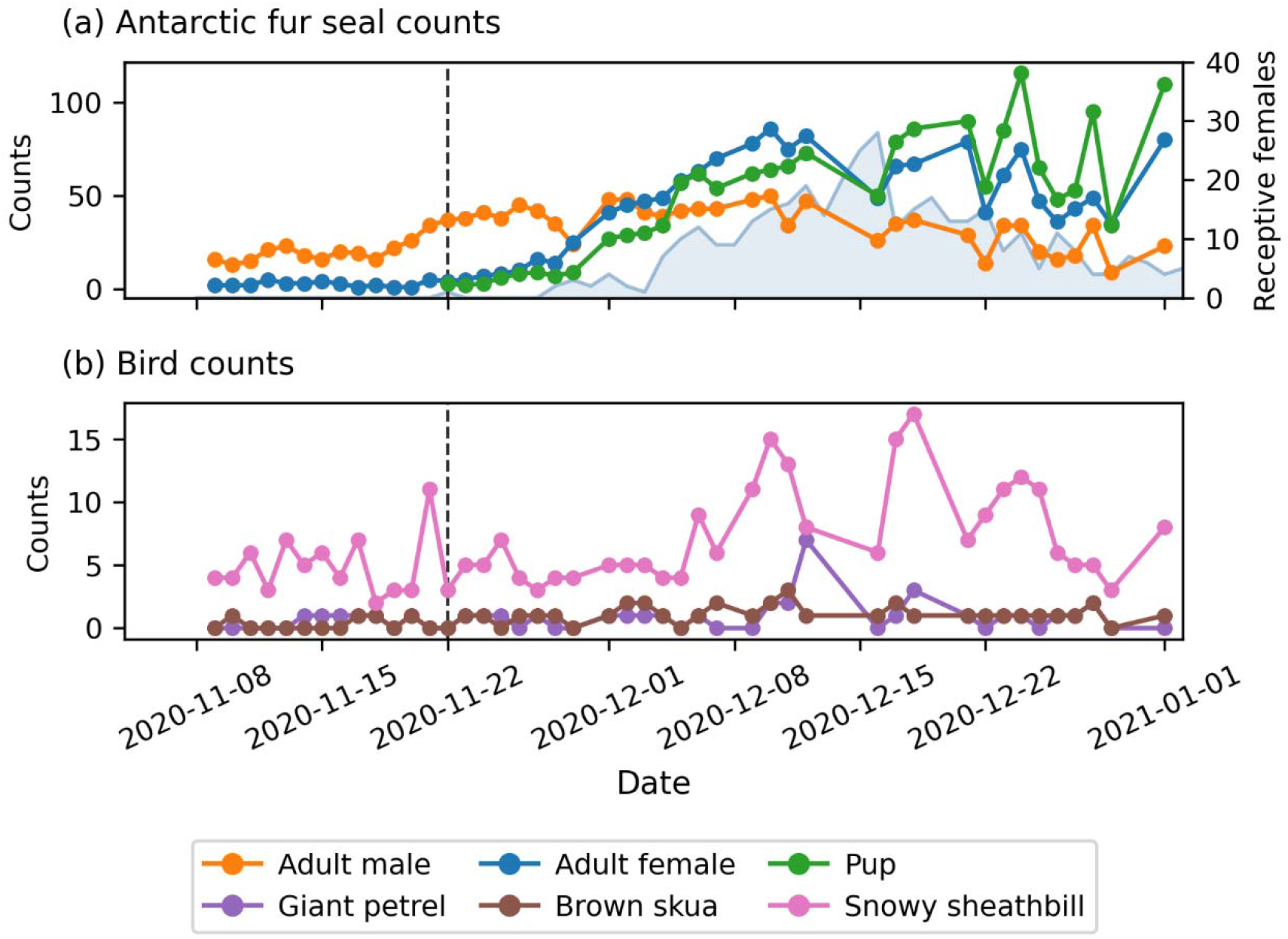
Temporal trends in abundance at the Special Study Beach. (a) Daily maximum counts obtained via the neural network of Antarctic fur seals classified as adult males (orange), adult females (blue), and pups (green). Light blue shading indicates the number of receptive adult females. (b) Daily maximum counts of giant petrels (purple), brown skuas (brown), and snowy sheathbills (pink). The dashed vertical black lines indicate the date of the first-born pup on SSB (22nd of November).

### (b) Colony-specific patterns in seal density

Next, to assess sex- and age-specific density patterns, we quantified seal density at FWB and SSB within the main breeding areas, defined by the 99% occupancy area of adult females and pups (Figure 3). Adult male densities were low at both colonies (FWB: 0.02 animals / m^2^; SSB: 0.04 animals / m^2^), with an approximately two-fold higher density at SSB. In contrast, adult female and pup densities differed substantially between colonies: females were more than five times denser at SSB (0.69 animals / m^2^) than at FWB (0.12 animals / m^2^), and pup densities followed the same pattern (FWB: 0.12 animals / m^2^; SSB: 0.50 animals / m^2^), reflecting a four-fold difference. Consequently, total seal density was four-fold higher at SSB (1.13 animals / m^2^) than at FWB (0.27 animals / m^2^), consistent with previously reported census-based estimates (FWB: 0.29 animals / m^2^; SSB: 1.13 animals / m^2^; Meise et al. 2016), suggesting that this density difference has remained stable for at least a decade.

**Figure 3:**
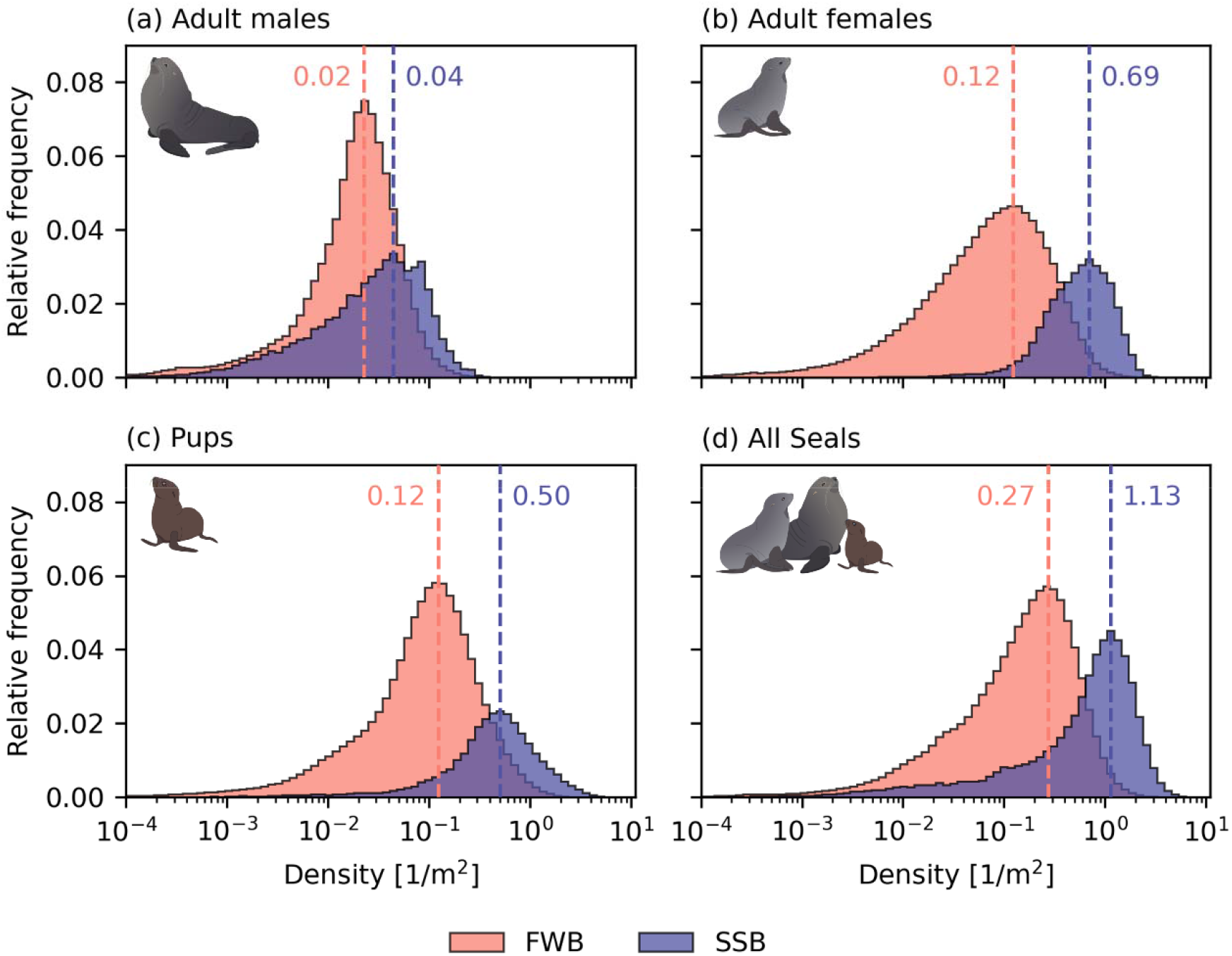
Colony specific patterns of seal density. Each panel compares the Voronoi density distribution of Antarctic fur seal categories and all seals combined: (a) adult males, (b) adult females, (c) pups and (d) all seals combined at FWB (distributions in salmon) and SSB (distributions in indigo). Dashed vertical lines indicate the peak values of the respective distributions, and the value is given next to the line. Density estimates for adult females and pups at SSB are corrected for scaffolding occlusion (see Methods). Original artwork by ALB.

As the scaffolding at SSB can occlude smaller animals, we compared corrected and uncorrected density estimates (Supplementary Figure 5), revealing a slight underestimation of females and a stronger underestimation of pups when scaffolding was not accounted for. But even the uncorrected densities indicate that the overall density difference between colonies is driven by a markedly higher density of females and their pups at SSB.

### (c) Abundance ratios of birds to pups between colonies

As direct measures of predation risk are difficult to obtain, we used bird-to-pup abundance ratios as a proxy, calculated for each species using a 7-day rolling window, to compare predation risk between FWB and SSB (Figure 4). At FWB, the median rolling bird-to-pup ratio was 0.19 (95% CI = 0.04– 1.15) for giant petrels and 0.44 (95% CI = 0.05–2.83) for brown skuas. At SSB, these ratios were significantly lower at 0.03 (95% CI = 0.01–0.19) for giant petrels and 0.03 (95% CI = 0.01–0.17) for brown skuas (both p < 0.001), suggesting that pups at SSB experience lower avian predation risk. The season-integrated ratios reinforced this pattern, with FWB showing approximately 4–8 times higher bird-to-pup ratios than SSB for both giant petrels (FWB = 0.09, SSB = 0.02) and brown skuas (FWB = 0.18, SSB = 0.02).

**Figure 4:**
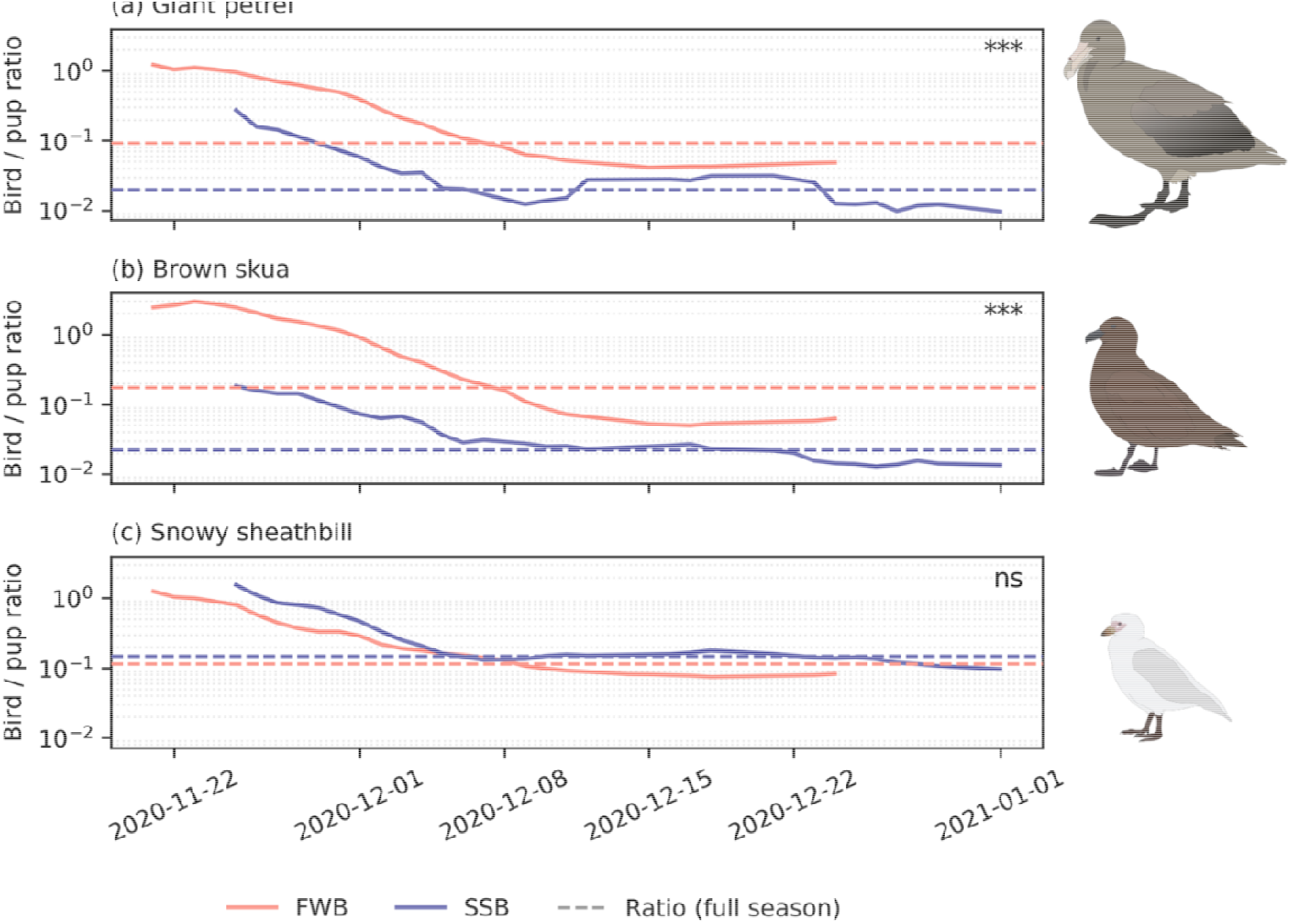
Abundance ratios of birds to pups across colonies. The panels show the bird-to-pup abundance ratios for (a) giant petrels, (b) brown skuas, and (c) snowy sheathbills across the pupping season. Solid lines (FWB = salmon, SSB = indigo) show 7-day rolling abundance ratios, whereas dashed lines represent the season-integrated ratios. Significant differences between colonies are indicated by asterisks: * p ≤ 0.05, ** p ≤ 0.01, *** p ≤ 0.001, ns = non-significant. Original artwork by ALB.

**Figure 5:**
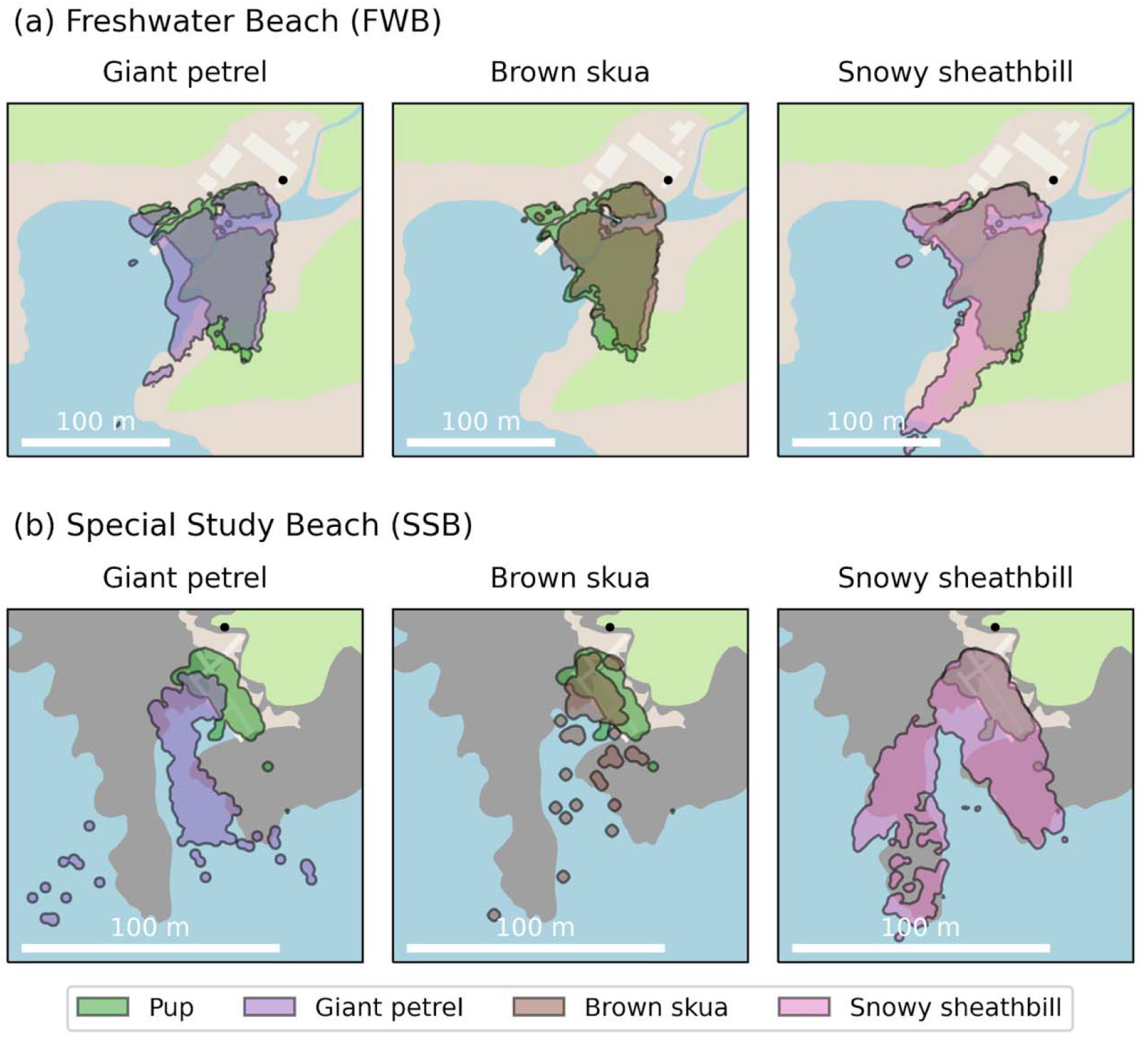
Spatial associations between birds and pups across colonies. Each panel visualises the overlap of the 99% kernel density estimates representing the spatial distributions of one of the three bird species (snowy sheathbill, brown skua, and giant petrel) and Antarctic fur seal pups at (a) FWB and (b) SSB. The black points indicate the camera positions.

These differences were most pronounced early in the pupping season, when pup numbers were still low. By contrast, the bird-to-pup ratio for snowy sheathbills did not differ significantly between colonies, with similar median rolling ratios (FWB = 0.18, 95% CI = 0.08–1.12; SSB = 0.16, 95% CI = 0.10–1.24; p = 0.90) and comparable season-integrated ratios (FWB = 0.12, SSB = 0.15), indicating that sheathbill presence scaled proportionally with pup numbers at both colonies.

### (d) Spatial associations between birds and pups

To test if higher density leads to the exclusion of avian predators from the main breeding colony, we calculated the proportion of the occupancy area of each bird species that overlapped with the pup occupancy area. At FWB, all three bird species showed extensive spatial overlap with the pups’ spatial distribution (giant petrel = 86%; brown skua = 79%; snowy sheathbill = 94%). At SSB, the overlap between giant petrels and pups was only 25% and was mainly limited to the shoreline, consistent with the drowning hunting strategy of giant petrels (Nagel *et al*. 2022). The spatial overlap between brown skuas and pups was moderate at 53%, whereas snowy sheathbills overlapped almost entirely with the pups (99%). This suggests that predators, but not scavengers, are physically excluded from SSB, most likely due to the much higher density of adult females.

## Discussion

This study investigated seal density and spatio-temporal predator-prey patterns at two Antarctic fur seal colonies of contrasting density at Bird Island, South Georgia. We employed automated camera systems together with a state-of-the-art object detection model to generate a high-resolution dataset of automatic detections. We used these data to characterise patterns in seal density, quantify colony differences in bird-to-pup abundance ratios and to compare spatial associations between pups and bird species. Together, our results shed light on how predator-prey associations are shaped by density.

### Training the neural network for a new location

To analyse the images recorded at SSB, we adopted the widely used YOLOv8 architecture (Jocher *et al*. 2023) rather than the custom hierarchical model used at FWB (Berthelsen & Bartl 2026), to improve reproducibility and accessibility. This lowers the technical barrier for future applications of this approach, as the network achieved high-quality detections comparable to those obtained at FWB. SSB posed additional detection challenges relative to FWB: whereas FWB is an open beach affording largely unobstructed camera views, SSB is partially occluded by elevated walkways and scaffolding infrastructure, complicating object detection, which most likely caused the underestimation of adult female and in particular pup abundance (Supplementary Figure 5). We therefore excluded the area obstructed by the scaffold when calculating corrected densities, which were then used for all subsequent analyses. Regardless of this correction, the model performed robustly under these conditions, reliably detected the seasonal abundance patterns of Antarctic fur seals, and thereby demonstrated the generalisability of this approach across colony settings of varying complexity.

### Temporal trends in abundance at SSB

Automated counts at SSB were validated against manual and census counts and closely resembled the seasonal abundance patterns observed at FWB (Berthelsen & Bartl 2026), consistent with the known breeding phenology of Antarctic fur seals(Duck 1990; McCann 1980). Importantly, we observed a nearly complete absence of giant petrels and brown skuas at SSB. Anecdotally, fewer giant petrels have been observed at SSB (Nagel *et al*. 2021), and their consistent numbers over the season at FWB suggest that intraspecific competition limits their numbers rather than prey abundance. The absence of brown skuas from SSB suggests that they forage elsewhere, or that the higher density of seals effectively excludes them, as has been observed for colonially breeding bird species (Busdieker *et al*. 2020; Lee *et al*. 2025). Snowy sheathbill abundance peaked during the height of the breeding season, when the beach is littered with placentae, carrion and faeces, reinforcing their role as scavengers (Burger 1981).

### Colony-specific patterns in seal density

To characterise colony-specific patterns in seal densities, we quantified local densities for each category and for all seals combined. Census counts obtained a decade ago revealed that the two colonies differ about four-fold in density (Meise *et al*. 2016); however this analysis did not explore differences between the sexes or life history stages. Adult male fur seals defend territories at considerable cost (Baker & McCann 1989; Boyd & Duck 1991), yet successful males reap the benefits by fathering the majority of the pups (Hoffman *et al*. 2003). As territory-holding males require a minimum amount of space, their densities are constrained, and the similarly low densities observed at both colonies (FWB: 0.02 animals / m^2^; SSB: 0.04 animals / m^2^) are consistent with this. The two-fold higher density at SSB likely reflects smaller territory sizes imposed by spatial constraints at SSB or increased competition, rather than a fundamentally different spacing behaviour. By contrast, both adult female and pup density differed strongly between colonies, potentially due to strong site fidelity among females (van Benthem *et al*. 2026; Hoffman & Forcada 2012). These pronounced differences in density (five-fold and four-fold difference in adult female and pups, respectively) are likely the primary driver of the overall four-fold density difference. The higher density of breeding females at SSB could indicate that the males holding territory there are of superior quality (van Benthem *et al*. 2026).

### Abundance ratios of birds to pups between colonies

To formally test for differences in bird abundance relative to pup abundance between colonies, we calculated bird-to-pup abundance ratios, which were significantly higher at FWB compared to SSB for giant petrels and brown skuas. This confirmed that the greater bird numbers at FWB reflected a genuinely higher predator presence rather than simply a larger pup population attracting more birds. Meanwhile, we found that the abundance ratios of snowy sheathbills to pups were comparable between colonies. These patterns align with the temporal associations observed at FWB, where snowy sheathbill abundance was positively associated with pup abundance, whereas giant petrel and brown skua abundances were unlinked to pup abundance (Berthelsen & Bartl 2026).

The higher ratios at FWB indicate that pups born at the low-density colony are more likely to encounter giant petrels and brown skuas, suggesting a higher predation risk. Taken together with the observation of elevated pup mortality at FWB due to predation (Nagel *et al*. 2021), this lends further support to a predator-driven Allee effect. However, as pup numbers increase over the course of the breeding season both colonies experienced declines in bird-to-pup abundance ratios, which could indicate that pups overall benefit from safety in numbers, also known as the dilution effect (Bednekoff & Lima 1998). Concurrently, the similarity in snowy sheathbill to pup ratios on FWB and SSB suggests a comparable rate of scavenging between colonies.

### Spatial associations between birds and pups

We compared spatial associations between Antarctic fur seal pups and the predator-scavenger bird species at FWB and SSB by estimating the overlap between bird and pup 99% occupancy areas. Assuming that spatial overlap serves as a proxy for predation risk and scavenging pressure, the extensive overlap observed at FWB between all three focal bird species and pups indicates that both predation and scavenging are likely prevalent there. Comparatively, a gradient of spatial overlap from low for giant petrels through intermediate for brown skuas to high for snowy sheathbills was found at SSB, suggesting that scavenging is commonplace, but predation risk is low.

At FWB, only giant petrels were extensively found in the tidal zone, brown skuas closely matched the spatial distribution of adult females and their pups, whereas snowy sheathbills occupied the entire colony. These spatial patterns of occupancy likely arise from foraging niche partitioning (Berthelsen & Bartl 2026). While the spatial distributions of the three focal bird species on SSB broadly reflect the same patterns, the reduced overlap suggests that the high density of adult female Antarctic fur seals might constrain predator access, thereby reducing hunting opportunities within the colony. Given that the occupancy area of giant petrels clearly follows the channel, tidal variation likely affected how far up the beach this species was observed. As the spatial overlap was aggregated across the entire season, the actual spatial overlap between giant petrels and pups might be even narrower than reported here. However, as drowning is one of the main predation strategies of the giant petrels, their restriction to the shoreline may reflect an adaptation to capitalise on this strategy (Nagel *et al*. 2022). Taken together, these patterns suggest that adult females are highly tolerant of snowy sheathbills, likely due to their role as scavengers (Burger 1981), intermediate tolerance for brown skuas and low tolerance for giant petrels, reflecting the differing levels of threat these species pose to pups.

### Caveats

A few caveats should be considered when interpreting our findings. We used spatial proximity as a proxy for predation risk and scavenging pressure rather than directly observing predation events. However, the one-minute resolution of our automated camera system precludes the direct capture of predation events, which could be improved by higher temporal resolution. In addition, camera field-of-view constraints at SSB — where scaffolding and walkways partially obstruct the view — likely caused us to underestimate the abundance of adult females and particularly pups, as their smaller body size makes them more likely to be completely obscured compared to adult males. However, these obstructions did not systematically bias the bird-to-pup ratios as birds and pups are of comparable size and therefore equally likely to be obscured.

### Perspectives

Our results suggest that adult females may be important deterrents of predators. To explore this further, one would need to distinguish whether predator exclusion from the main breeding area is a passive or active process. This could be experimentally tested by presenting adult females with models representing each of the focal bird species, or simply bird models representing a size gradient, to quantify whether females respond differently to each of them. Furthermore, climate-driven reductions in food availability (Forcada *et al*. 2023; Forcada & Hoffman 2014) will most likely influence these dynamics. As food becomes scarce, adult females spend longer foraging at sea (Nagel *et al*. 2022), increasing pup predation risk. Should this persist, a general decline in colony density might cause the density at SSB to fall below a critical threshold where it is no longer high enough to effectively exclude predators. A predator-driven Allee effect has already been documented at FWB, indicating that such a threshold exists. Thus, continuous monitoring at SSB or across a range of colonies differing in density could narrow down the estimate of this threshold and provide predictive insights into the long-term effects of climate change on predator-prey dynamics.

### Conclusion

In summary, this study demonstrates the value of using autonomous time-lapse cameras in combination with neural network-based analysis for investigating predator-prey interactions in a colonial breeder. This approach provides comprehensive, near-continuous observations of temporal and spatial abundance patterns while minimizing human disturbance. Comparison of patterns of density in two Antarctic fur seal colonies of contrasting seal density suggested that the density difference between the colonies has persisted for at least a decade, largely driven by a higher density of adult females and their pups. Furthermore, our results revealed lower predation risk for pups in the high-density colony, primarily driven by the exclusion of predatory birds from the central breeding area, whereas the scavenger birds were consistently present within the colony. Together, these findings support a predator-driven Allee effect, whereby higher local density reduces predation risk.

## Funding Statement

J.B. DFG (German Research Foundation) priority programme “Antarctic Research with Comparative Investigations in Arctic Ice Areas” (SPP 1158, project number ZI1527/7-1)

A.L.B. Collaborative Research Centre Transregio 212 “A Novel Synthesis of Individualisation across Behaviour, Ecology and Evolution: Niche Choice, Niche Conformance, Niche Construction (NC3)” (SFB TRR 212, project numbers 316099922 & 396774617)

A.W. DFG (German Research Foundation) priority programme “Antarctic Research with Comparative Investigations in Arctic Ice Areas” (SPP 1158, project number 443134677) C.F.-C. The core science programme of the British Antarctic Survey, Polar Science For a Sustainable Planet, NERC-UKRI (Natural Environment Research Council, United Kingdom UKRI).

J.F. The core science programme of the British Antarctic Survey, Polar Science For a Sustainable Planet, NERC-UKRI (Natural Environment Research Council, United Kingdom UKRI).

R.N. Collaborative Research Centre Transregio 212 “A Novel Synthesis of Individualisation across Behaviour, Ecology and Evolution: Niche Choice, Niche Conformance, Niche Construction (NC3)” (SFB TRR 212, project numbers 316099922 & 396774617)

J.I.H. DFG (German Research Foundation) priority programme “Antarctic Research with Comparative Investigations in Arctic Ice Areas” SPP 1158 (project number 424119118). Collaborative Research Centre Transregio 212 “A Novel Synthesis of Individualisation across Behaviour, Ecology and Evolution: Niche Choice, Niche Conformance, Niche Construction (NC3)” (SFB TRR 212, project numbers 316099922 & 396774617).

B.F. DFG (German Research Foundation) priority programme “Antarctic Research with Comparative Investigations in Arctic Ice Areas” (SPP 1158, project number 443134677)

## Acknowledgments

The authors thank Iain Angus Gordon and David Reid for technical support in the field. The authors gratefully acknowledge the scientific support and HPC resources provided by the Erlangen National High Performance Computing Center (NHR@FAU) of the Friedrich-Alexander-Universität Erlangen-Nürnberg (FAU). Support for the article processing charge was granted by the DFG and the Open Access Publication Fund of Bielefeld University.

## Ethical Statement

The work at Bird Island was carried out by BAS under permits from the Government of South Georgia and the South Sandwich Islands (Wildlife and Protected Areas Ordinance (2011), RAP permit number 2020/015).

## Data Accessibility

The trained model, automated detection counts, annotated training images are available on Zenodo (DOI: 10.5281/zenodo.18955385). The code required to reproduce all analyses is available on GitHub (https://github.com/fabrylab/AntarcticFurSealDensity)

## Competing Interests

The authors declare no competing interests

## Authors’ Contributions

J.I.H., R.N. and B.F. conceived the study. J.I.H. acquired funding for the project. R.N. and C.F.-C. conducted the fieldwork and C.F.-C. maintained the camera. A.W. trained the neural network for FWB and build the camera system, J.B. trained the neural network for SSB. J.B. and A.L.B. performed the formal analyses and visualized the results. A.L.B., J.B., B.F. and J.I.H. wrote the original draft. All authors provided feedback and approved the final manuscript.

## Notes

https://github.com/fabrylab/AntarcticFurSealDensity

https://zenodo.org/records/18955385

### Competing Interest Statement

The authors have declared no competing interest.

