## Supplementary Material for "High population density limits predator access in Antarctic fur seal breeding colonies"

**Supplementary information**

**Neural network**

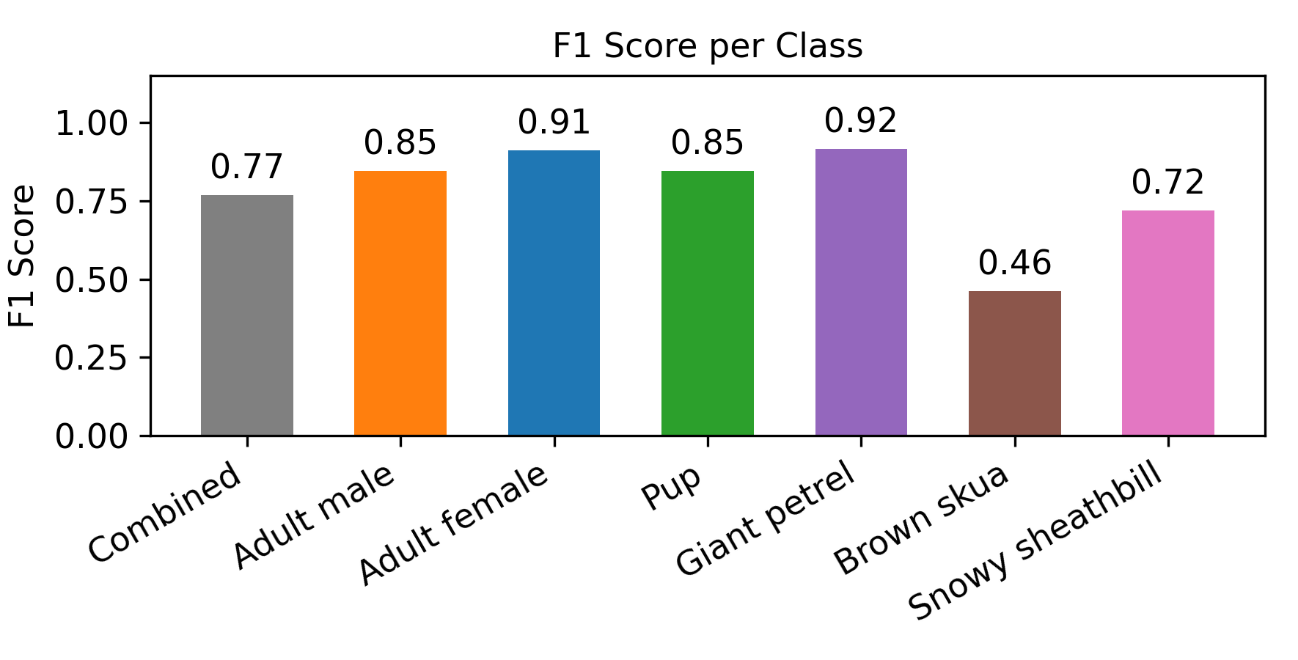

**Supplementary Figure 1: F1 scores of the object detection model for each class.** The neural network was trained on nine distinct classes, of which six focal classes are shown: the three Antarctic fur seal categories (adult male, adult female, pup) and three associated bird species (giant petrel, brown skua, snowy sheathbill). The mean F1 score across all six classes is shown in grey. F1 scores were evaluated at a confidence threshold of 0.65.

The brown skua shows the lowest F1 score (0.46), likely reflecting both its relative rarity in the training data and its visual similarity to fur seal pups in colouration, making it the most challenging class for the model to distinguish. However, comparison with manual counts (Supplementary Figure 3) shows that the model tends to overestimate rather than underestimate skua abundance, indicating that the lower F1 score reflects occasional false positives rather than missed detections. As overestimation is conservative with respect to our findings, with any bias working against detecting differences between colonies, the results reported in our manuscript remain valid.

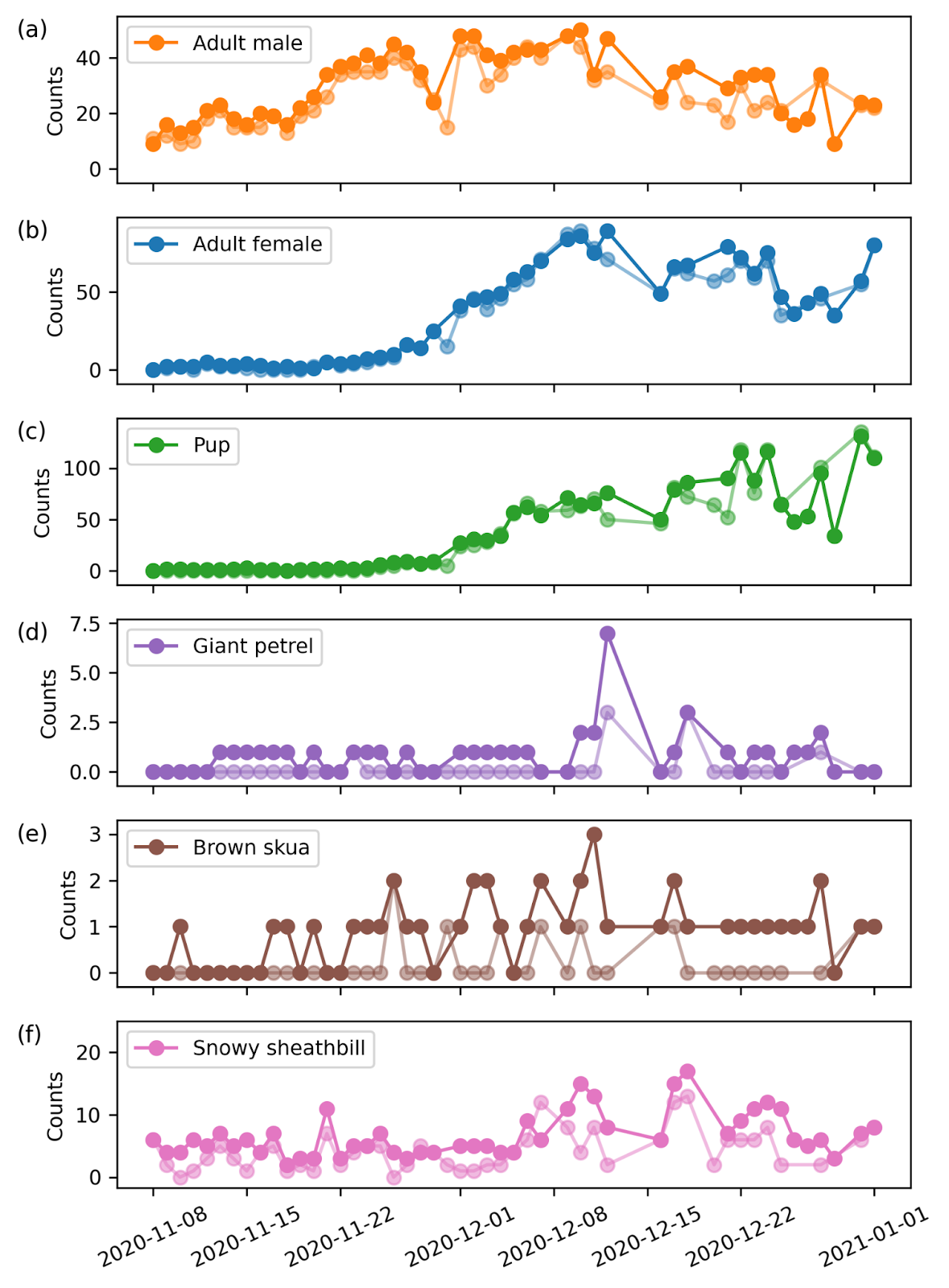

**Supplementary Figure 2: Comparison of manual and automated counts at the Special Study Beach.** Daily maximum counts from a neural network automatic detection algorithm (solid colours) compared with manual counts (semi-transparent colours) for Antarctic fur seals: (a) adult males, (b) adult females and (c) pups, as well as the predator species: (d) giant petrels, (e) brown skuas and (f) snowy sheathbills.

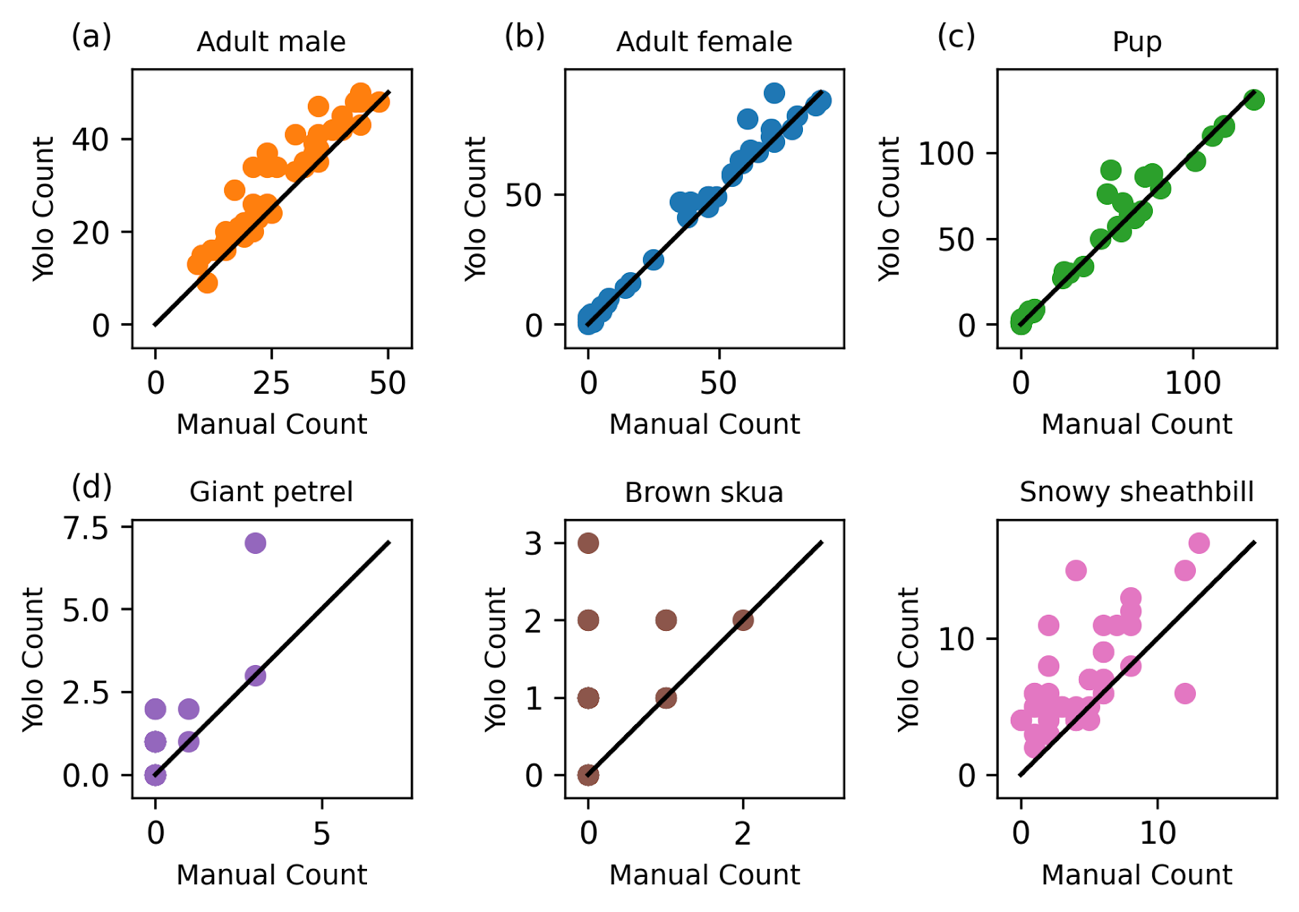

**Supplementary Figure 3: Agreement between automated and manual counts at the Special Study Beach.** Comparison of daily maximum neural network counts against manual counts for each focal class: Antarctic fur seal categories (a) adult male, (b) adult female, and (c) pup, and associated bird species (d) giant petrel, (e) brown skua, and (f) snowy sheathbill. The black diagonal line indicates perfect one-to-one agreement between the two counting methods.

**Extended methods**

Census counts are obtained by recording daily abundance of territorial males, breeding and non-breeding females, newborn pups and newly found dead pups within a designated area of the breeding beach. Non-territorial males occupy the rocky ridges surrounding the cobblestone beach, which fall within the camera FOV but outside the census area. During the afternoon, abundance counts are carried out for adult females, which mostly occupy the census area. Meanwhile, the birth and death of pups are recorded continuously throughout the day, and so the daily count of pups represent the cumulative count of born pups minus the number of dead pups, and not the number of pups currently present within the census area.

To compare the automated counts with census counts, we extracted the abundance from an area within the images, which correspond to the census area (Supplementary Figure 4). We obtained a near-perfect agreement for adult males, a slight underestimate of adult females, and a substantial disagreement for pups as the season progressed.

The discrepancy between automated and census counts for adult females and pups likely reflects the presence of the scaffolding, which obstructs parts of the census area from the image FOV. The impact of this obstruction scales inversely with body size: adult males are least affected, as their large size means some portion of the animal typically remains visible even when partially occluded. Adult females, and especially pups, are more likely to be entirely hidden behind obstructions, increasing the probability of missed counts relative to the census data. However, given the nature of the pup census counts the discrepancy could also reflect pups moving outside the camera FOV, seeking refuge in the surrounding tussock grass, which is not visible in the images. This interpretation is supported by the strong agreement previously confirmed between manual and automated pup counts within the visible area, suggesting the deficit arises from animals leaving the field of view rather than from detection errors.

Due to their smaller body size, adult females and particularly pups are more susceptible to complete occlusion behind the scaffolding and walkways at SSB, likely resulting in an underestimate of their abundance relative to the larger adult males, and consequently lower density estimates. Adult males were excluded from this correction for two reasons: first, census counts showed near-perfect agreement with automated male counts (Supplementary Figure 4), indicating that occlusion has negligible impact on male detection; second, due to their low density and large territory sizes, male Voronoi cells almost universally intersect the scaffolding area, making the correction uninformative. Importantly, even without any correction — using all detections regardless of proximity to the scaffolding — the density distributions at SSB already exceed those at FWB (indigo vs. salmon distributions), confirming that the colony density differences reported in the main analysis are robust and do not depend on the scaffolding correction. To further quantify the effect of occlusion, we calculated corrected density distributions by first outlining the scaffolding structure that blocks the field of view and projecting this outline into the top-view. Voronoi cells overlapping with the scaffolding area were then excluded, so that only individuals in unobstructed areas contribute to the corrected estimates (Supplementary Figure 5). The corrected peak densities (†) are shown for reference, illustrating that while the correction shifts the SSB distributions towards higher densities — consistent with a systematic underestimation of females and especially pups — the overall pattern of higher density at SSB holds regardless of whether the correction is applied.

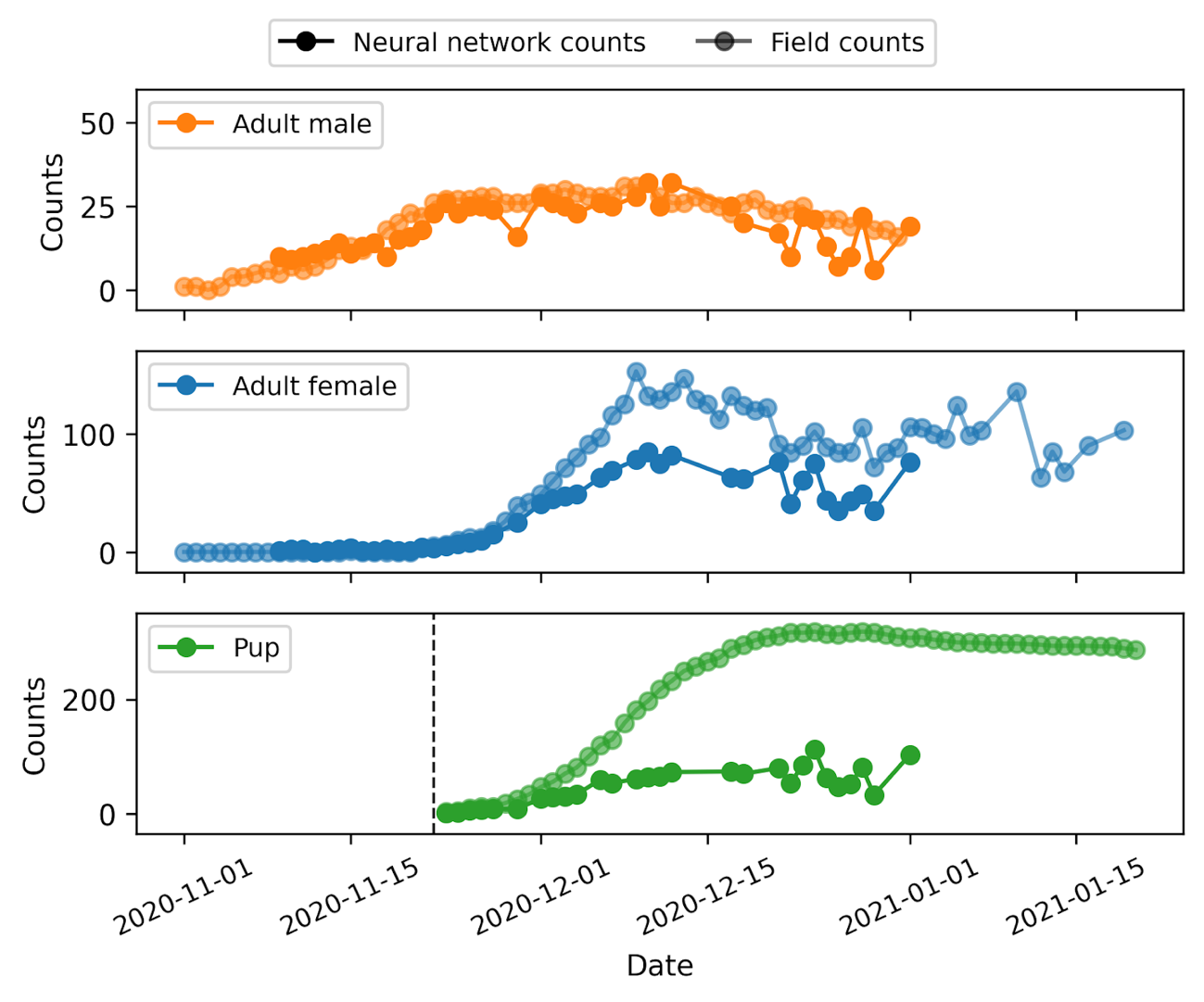
**Supplementary Figure 4: Comparison between automated and census counts at Special Study Beach.** Daily maximum counts from a neural network automatic detection algorithm (solid colours) compared with census counts (semi-transparent colours) for Antarctic fur seals: (a) adult males, (b) adult females and (c) pups.

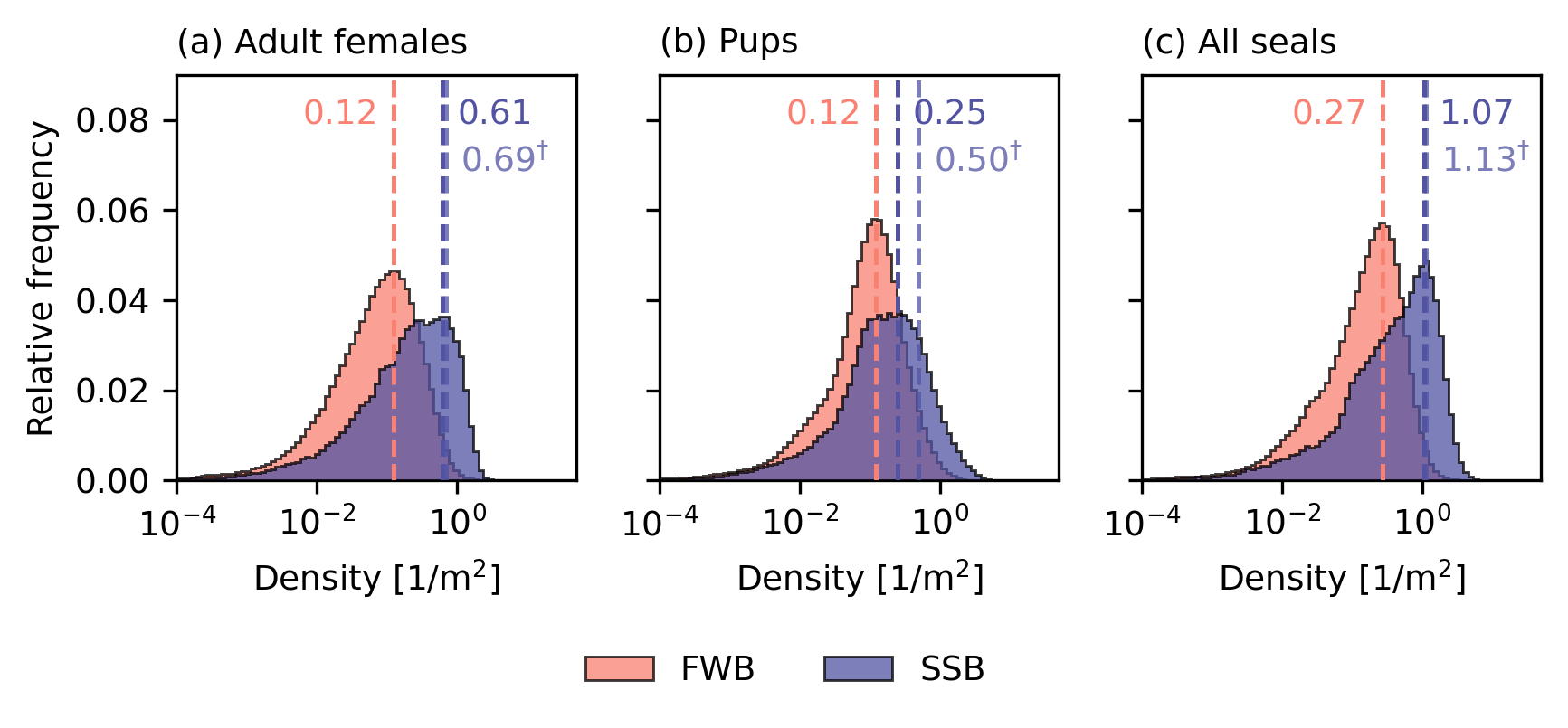

**Supplementary Figure 5: Effect of scaffolding occlusion on seal density estimates at Special Study Beach.** Panels show the Voronoi density distributions for (a) adult females, (b) pups, and (c) all seals combined, using all detected individuals regardless of their proximity to the scaffolding structure (i.e. the uncorrected estimates), for FWB (salmon) and SSB (indigo). Dashed vertical lines indicate the peak of each distribution; the corrected peak values (from Figure 3) are additionally shown as dagger-marked lines (†) for comparison. Adult males are not shown as their large body size renders them largely unaffected by scaffolding occlusion (see Supplementary Figure 4).
